# Gene model for the ortholog of *Ilp4* in *Drosophila simulans*

**DOI:** 10.64898/2026.02.06.704405

**Authors:** Leon F. Laskowski, Madeline L. Gruys, Robyn Huber, AnnaMaria DiGeronimo, Andrew M. Arsham, Vidya Chandrasekaran, Chinmay P. Rele, Lori Boies

## Abstract

Gene Model for *Insulin-like peptide 4* (*Ilp4)* in the *D. simulans* DsimGB2 assembly (GCA_000754195.3). The characterization of this ortholog was carried out as part of a larger, ongoing dataset designed to explore the evolution of the insulin/insulin-like growth factor signaling (IIS) pathway across the genus *Drosophila*, utilizing the Genomics Education Partnership gene annotation protocol within Course-based Undergraduate Research Experiences.

## Introduction

> This article reports a predicted gene model generated by undergraduate work using a structured gene model annotation protocol defined by the Genomics Education Partnership (GEP; thegep.org) for Course-based Undergraduate Research Experience (CURE). The following information in quotes may be repeated in other articles submitted by participants using the same GEP CURE protocol for annotating Drosophila species orthologs of Drosophila melanogaster genes in the insulin signaling pathway.

The insulin signaling pathway is a highly conserved regulatory system that plays a central role in controlling growth, metabolism, and development across metazoans (Biglou, *et* .*al*., 2021). In *Drosophila*, insulin signaling is mediated by a family of eight insulin-like peptides (Ilps) that act as ligands for the insulin receptor (Sharma *et*.*al*.,2019 & Okamoto & Yamanaka, 2015). Although these peptides share structural similarities, they differ in their spatial and temporal expression patterns and in their physiological roles. *Insulin-like peptide 4* (*Ilp4*) is expressed primarily in the embryonic mesoderm and midgut, yet despite this defined expression pattern and its strong evolutionary conservation, the specific biological functions of *Ilp4* remain poorly characterized (Brogiolo, *et. al*, 2001).

Comparative genomic studies have shown that among the insulin-like peptides *Ilp1* through *Ilp7, Ilp4* is one of the most highly conserved across *Drosophila* species, second only to *Ilp7* (Grönke, *et. al*., 2010). This conservation suggests functional importance, even as direct experimental evidence describing *Ilp4* activity remains limited. As a result, accurate annotation of *Ilp4* orthologs in multiple *Drosophila* species provides a necessary foundation for future functional and evolutionary analyses of this gene family.

*Drosophila simulans* is a close relative of *Drosophila melanogaster* and serves as an important comparative species for examining gene structure conservation, sequence divergence, and syntenic relationships. Its relatively recent divergence from *D. melanogaster* allows for detailed comparisons that can distinguish lineage-specific changes from conserved features of insulin signaling genes (Laskowski, *et. al*., . The gene model presented here represents the ortholog of *Ilp4* in the *D. simulans* genome and refines existing computational predictions through manual annotation supported by multiple lines of genomic evidence.

“In this GEP CURE protocol students use web-based tools to manually annotate genes in non-model Drosophila species based on orthology to genes in the well-annotated model organism, the fruit fly Drosophila melanogaster. This allows undergraduates to participate in course-based research by generating manual annotations of genes in non-model species (Rele et al., 2023).

Computational-based gene predictions in any organism are often improved by careful manual annotation and curation, allowing for more accurate analyses of gene and genome evolution (Mudge and Harrow, 2016; Tello-Ruiz et al., 2019). These models of orthologous genes across species, such as the one presented here, then provide a reliable basis for further evolutionary genomic analyses when made available to the scientific community.” (Myers et al., 2024).

## Results

### Synteny

The *Ilp4* ortholog is located on chromosome 3L in *D. melanogaster* and surrounded by *Ilp5* and *CG43987* (upstream) and *Ilp3* and *Ilp2* (downstream). Additionally, *CG32052* was found to be nesting *Ilp4, Ilp3*, and *Ilp2* within its first intron; *CG43987* was found to be nesting *Ilp5. Tblastn* showed the *Ilp4* (*LOC6737493)* ortholog was located on scaffold CM002912 in *D. simulans. Ilp4* (*LOC6737493*) is surrounded upstream by *LOC6737496* and *LOC6737495* (*Ilp5* and *CG43987*, respectively) and downstream by *LOC6747492* and *LOC6737491* (*Ilp3* and *Ilp2*, respectively). *CG32052* (*LOC6737494*) and *CG43987* (*LOC6737495*) were found to be nesting genes in *D. simulans*. The synteny diagram (Figure 1A) shows the genomic neighborhood in both *D. melanogaster* and *D. simulans*. Because the genomic neighborhood matches in both species, a low E-value and high percent identities were given by *blast*, we concluded that this region contains the ortholog for *Ilp4* in *D. simulans*.

**Figure 1:**
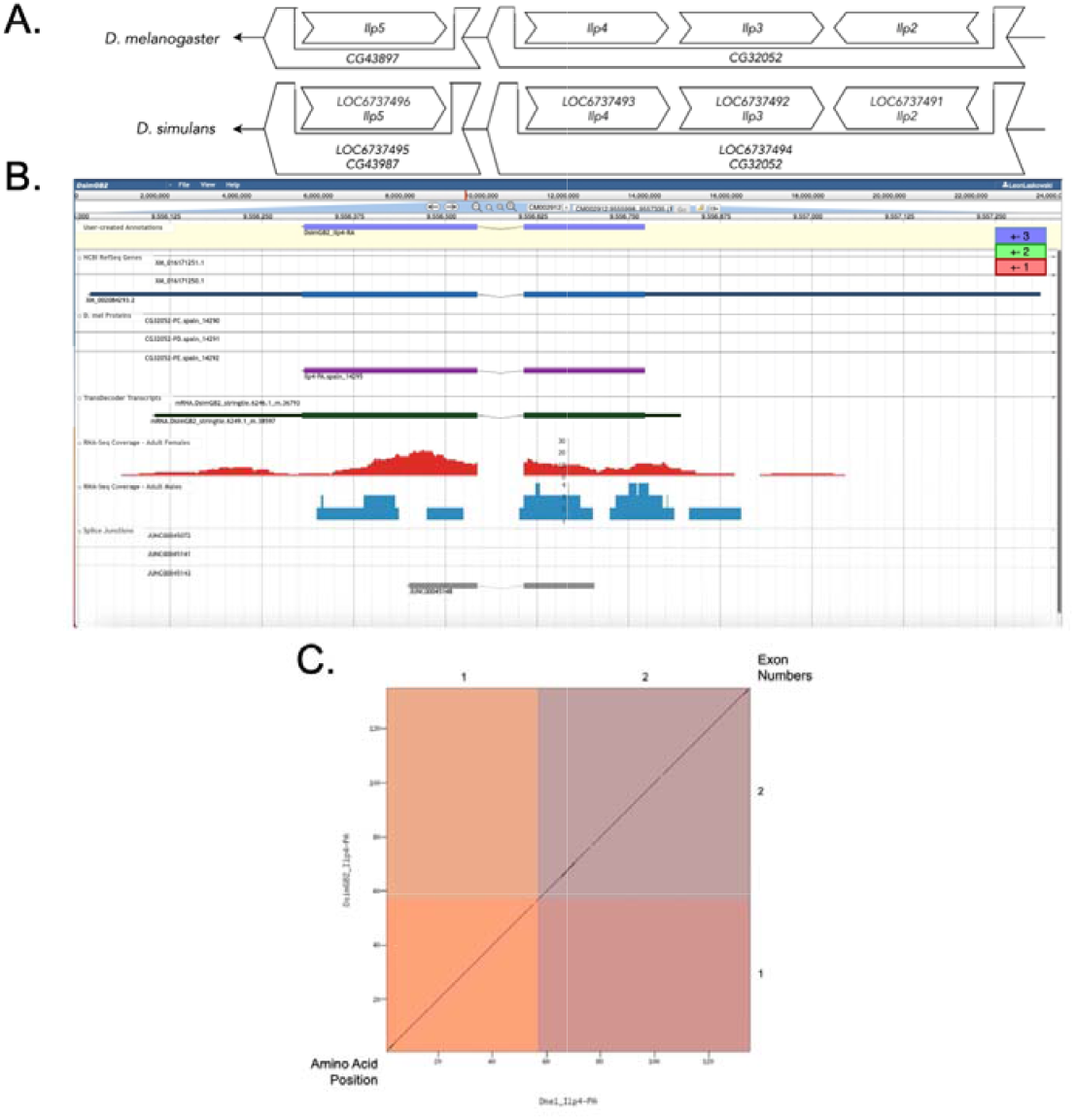
*Ilp4* gene model comparison between *Drosophila simulans* and *Drosophila melanogaster*. **(A) Synteny of genomic neighborhood of *Ilp4* in both *D. melanogaster* as well as *D. simulans***. The direction of the arrow at the end of the diagram for both *D. melanogaster* and *D. simulans* indicates the strand that the target gene resides on. The direction of the boxes in both species indicates the genes orientation relative to the target gene. The two bigger boxes surrounding smaller boxes represent nesting genes. **(B) Gene Model in Apollo:** A screenshot of the Apollo instance housing the gene model, NCBI RefSeq Genes, D. mel proteins, TransDecoder Transcript, RNA-Seq tracks for Adult Females and Adult Males (Graveley *et al*., 2011, SRP006203), and Splice Junctions, exon reading frames are indicated in blue, green, and red as in legend. Numbers on the bottom and left correspond to amino acid position, numbers on the top and right correspond to exon number. **(C) Dot Plot of Ilp4-PA gene model in D. melanogaster (x-axis) vs. the Ilp4-PA gene model in D. simulans (y-axis)**.

### Gene Model

*Ilp4* has only one identified isoform (Ilp4-PA) annotated in *D. melanogaster*. Ilp4-PA was identified in *D*. simulans and has two coding exons, which is consistent with the number of coding exons annotated for *Ilp4* in *D. melanogaster*. The Dot Plot (Figure 1C) shows that the protein sequence of *Ilp4* in *D. simulans* aligns with that of *Ilp4* in *D. melanogaster* and shows a high percent identity between both species.

## Methods

“Detailed methods including algorithms, database versions, and citations for the complete annotation process can be found in Rele et al. (2023). Briefly, students use the GEP instance of the UCSC Genome Browser v.435 (https://gander.wustl.edu; Kent WJ et al., 2002; Navarro Gonzalez et al., 2021) to examine the genomic neighborhood of their reference IIS gene in the *D. melanogaster* genome assembly (Aug. 2014; BDGP Release 6 + ISO1 MT/dm6). Students then retrieve the protein sequence for the *D. melanogaster* reference gene for a given isoform and run it using *tblastn* against their target *Drosophila* species genome assembly on the NCBI BLAST server (https://blast.ncbi.nlm.nih.gov/Blast.cgi; Altschul et al., 1990) to identify potential orthologs. To validate the potential ortholog, students compare the local genomic neighborhood of their potential ortholog with the genomic neighborhood of their reference gene in *D. melanogaster*. This local synteny analysis includes at minimum the two upstream and downstream genes relative to their putative ortholog. They also explore other sets of genomic evidence using multiple alignment tracks in the Genome Browser, including BLAT alignments of RefSeq Genes, Spaln alignment of *D. melanogaster* proteins, multiple gene prediction tracks (e.g., GeMoMa, Geneid, Augustus), and modENCODE RNA-Seq from the target species. Detailed explanation of how these lines of genomic evidenced are leveraged by students in gene model development are described in Rele et al. (2023). Genomic structure information (e.g., CDSs, intron-exon number and boundaries, number of isoforms) for the *D. melanogaster* reference gene is retrieved through the Gene Record Finder (https://gander.wustl.edu/~wilson/dmelgenerecord/index.html; Rele et al., 2023). Approximate splice sites within the target gene are determined using *tblastn* using the CDSs from the *D. melanogaste*r reference gene. Coordinates of CDSs are then refined by examining aligned modENCODE RNA-Seq data, and by applying paradigms of molecular biology such as identifying canonical splice site sequences and ensuring the maintenance of an open reading frame across hypothesized splice sites. Students then confirm the biological validity of their target gene model using the Gene Model Checker (https://gander.wustl.edu/~wilson/dmelgenerecord/index.html; Rele et al., 2023), which compares the structure and translated sequence from their hypothesized target gene model against the *D. melanogaster* reference gene model. At least two independent models for a gene are generated by students under mentorship of their faculty course instructors. Those models are then reconciled by a third independent researcher mentored by the project leaders to produce the final model. Note: comparison of 5’ and 3’ UTR sequence information is not included in this GEP CURE protocol.” (Gruys et al, 2025)

## Supporting information

FASTA, GFF, PEP Files

## Supplemental Material

1. Zip file containing FASTA, PEP, GFF files for the gene model
2. Figure 1 in high resolution

**Metadata:** Bioinformatics, Genomics, Drosophila, Genotype Data, New Finding

## Acknowledgements

We thank Wilson Leung (Washington University, St. Louis) for developing and maintaining the technological infrastructure that supported the creation of this gene model, as well as Chinmay Rele and Laura K. Reed (University of Alabama) for their guidance and encouragement throughout the project. We are also grateful to FlyBase for providing the authoritative database for *Drosophila melanogaster* gene models. FlyBase is supported by grants NHGRI U41HG000739 and U24HG010859, UK Medical Research Council MR/W024233/1, NSF 2035515 and 2039324, BBSRC BB/T014008/1, and Wellcome Trust PLM13398.

## Funding

This material is based upon work supported by the National Science Foundation (1915544) and the National Institute of General Medical Sciences of the National Institutes of Health (R25GM130517) to the Genomics Education Partnership (GEP; https://thegep.org/; PI-Laura K. Reed). Any opinions, findings, and conclusions or recommendations expressed in this material are solely those of the author(s) and do not necessarily reflect the official views of the National Science Foundation nor the National Institutes of Health.

## Notes

### Competing Interest Statement

The authors have declared no competing interest.

